# Human-specific changes in two functional enhancers of *FOXP2*

**DOI:** 10.1101/157016

**Authors:** Antonio Benítez-Burraco, Raúl Torres-Ruiz, Pere Gelabert Xirinachs, Carles Lalueza-Fox, Sandra Rodríguez-Perales, Paloma García-Bellido

## Abstract

Two functional enhancers of *FOXP2,* a gene important for language development and evolution, exhibit several human-specific changes compared to extinct hominins that are located within the binding site for different transcription factors. Specifically, Neanderthals and Denisovans bear the ancestral allele in one position within the binding site for SMARCC1, involved in brain development and vitamin D metabolism. This change might have resulted in a different pattern of *FOXP2* expression in our species compared to extinct hominins.

## Main text

Mutations in the coding region of *FOXP2* are known to cause speech and language impairment (Watkins et al. 2002; Shriberg et al. 2006; Reuter et al. 2017). Modern humans and extinct hominins (Neanderthals and Denisovans) are thought to be endowed with distinctive language abilities, but no differences in the coding region of *FOXP2* have been found between species (Benítez-Burraco et al. 2008; Johansson 2015). Changes in the expression pattern/level of *FOXP2* have been thus hypothesised to contribute to the species-specific idiosynchratic language profile. Interestingly, Neanderthals bear the ancestral allele of a binding site for the transcription factor POU3F2 within intron 8 of *FOXP2,* which is more efficient in activating the transcription of the target gene (Maricic et al. 2013). Accordingly, higher levels of FOXP2 might be expected for this hominin species. In humans, partial duplications of *FOXP2* are related to delayed speech and language development (nssv13656147) and autism features with intellectual disability (nssv13648704).

We have recently uncovered two enhancers of *FOXP2* in the intergenic region between *FOXP2* and its adjacent *MDFIC* gene and validated their functionality by CRISPR/Cas9 (Torres-Ruiz et al. 2016). Deletion of any of them in the SK-N-MC neuroblastoma cell line downregulates *FOXP2* and decreases FOXP2 protein levels. This suggests that both elements upregulate *FOXP2* in vivo in brain cells. In this letter, we describe several differences in the sequences of these two enhancers between Neanderthals, Denisovans, and modern humans, which support the view that the expression of *FOXP2* may be differentially regulated in these three hominin species, plausibly impacting on the sensorimotor loops for auditory-vocal control to which FOXP2 contributes.

In doing so, we first compared the sequence of the two enhancers in several species of interest, namely, vertebrates in which imitative vocalization has been extensively studied in the context of language evolution. These include songbirds (zebra finch, collared flycatcher, white-throated sparrow, medium ground finch), cetaceans (dolphins, killer whales), bats (black flying-foxes, megabats, David’s myotis, microbats, big brown bats), primates (humans, chimps, gorillas, orangutans, gibbons, rhesus, baboons, marmosets, bushbabies), and rodents (mice). We then aligned the Denisovan and Neanderthal sequences with the human reference genome (GRCh37), looking for differences in the sequences of the two enhancers. Finally, we interrogated in silico whether the observed differences affect the regulatory properties of the two enhancers, and ultimately, *FOXP2* expression.

We found that these two enhancers are absent in songbirds (Figure 1). The proximal one (FOXP2-E^proximal^) is found in primates, bats and cetaceans, but not in rodents, whereas the distal enhancer (FOXP2-E^distal^) is partially conserved in cetaceans, some bats, and most primates (Figure 1). Interestingly, most of the predicted binding sites for transcription factors within the two enhancers are absent in the primate species that are most distantly related to humans (bushbabies and marmosets) (Figure 1). Regarding extinct hominins, in spite of the high degree of homology with modern humans, we found some differences between the human reference genome (GRCh37) and the genomic sequences from Neanderthals and Denisovans. The Denisovan FOXP2-E^proximal^ exhibits six differences with the modern human sequence, with two of these changes (highlighted in blue in Figure 1) located within predicted targets for transcription factors. The Neanderthal FOXP2-E^proximal^ shows five differences with the human enhancer, with two of them (highlighted in red in Figure 1) located within the predicted binding sites for the transcription factors POLR2A and SMARCC1. Only one change is shared by both species (114460158T>G, hg19), which affects the binding site for SMARCC1. Regarding FOXP2-E^distal^, there are no differences between the Denisovan sequence and the human sequence, whereas we found two differences with the Neanderthal enhancer, although they are located outside any predicted binding site for transcription factors (Figure 1).

**Figure 1.**
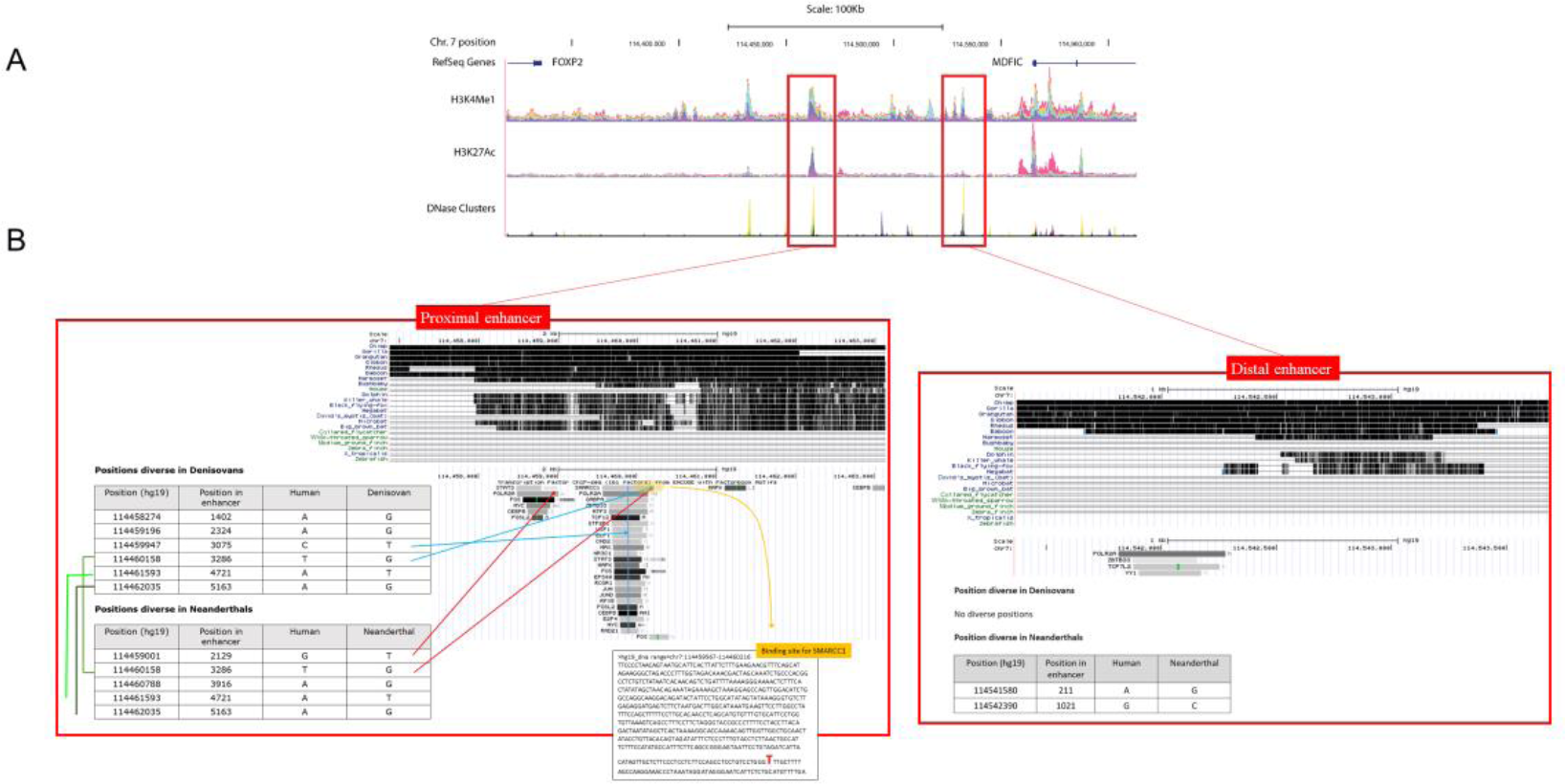
Evolutionary changes in the two *FOXP2* enhancers. A. Snapshot of an ENCODE UCSC genome-browser search showing the genomic location of the two enhancers (red boxes), downstream *FOXP2* and upstream *MDFIC,* and of three common hallmarks of cis-regulatory elements of gene expression: H3K4Me1 marks, H3K27Ac marks, and DNAse clusters. B. Changes in the sequences of the two enhancers in species of interest for language evolution, from songbirds to extinct hominins. Main evolutionary findings are shown inside the two red boxes, one for FOXP2-E^proximal^ (left) and one for FOXP2-E^distal^ (right). For each enhancer, we show the results of the MultiZ alignment of the sequences from selected extant vertebrates (up) and the differences with the Neanderthal and Denisovan sequences (bottom). The tables (bottom left) report the position and nature of the nucleotide differences between the human and the extinct hominin sequences. The transcription factor tracks (bottom right) show the transcription factor binding sites that are found within the enhancers (from a collection of ChIP-seq experiments). The darkness of the bar is proportional to the maximum signal strength. For SMARCC1, we provide the full binding sequence within FOXP2-E^proximal^, in which the derived T has been highlighted in red (box at the bottom).

*FOXP2* is a gene important for speech and language development and evolution (Scharff and Petri 2011; Graham and Fisher 2015). In view of the identity observed at the protein level between the three hominin species, we regard this nucleotide difference in the binding site for SMARCC1 within FOXP2-E^proximal^ of interest for inferring evolutionary modifications of *FOXP2* expression in the human clade. As noted, both abnormally low and abnormally high levels of FOXP2 protein result in abnormal cognitive phenotypes, which entail problems with speech and language. SMARCC1, a component of the large ATP-dependent chromatin remodelling complex SNF/SWI, plays an important role in the development of the forebrain, particularly, in neurogenesis (Narayanan et al. 2015). The implicated regulatory mechanism also involves *PAX6* (Ninkovic et al. 2013), which controls *FOXP2* expression (Coutinho et al. 2011). It has been hypothesised that selected differences in the coding and/or regulatory regions of *PAX6* and some of its partners might account for some of the cognitive differences between Neanderthals and humans (see Benítez-Burraco and Boeckx, 2015 for discussion). Interestingly, one of these partners is *SPAG5,* which helps PAX6 to regulate neuronal proliferation (Asami et al. 2011) and which exhibits several fixed changes in modern humans (Green et al. 2010; Meyer et al. 2012; Prüfer et al. 2013).

SMARCC1 is also involved in vitamin D-coupled transcription regulation (Hawes et al. 2015). Vitamin D deficiency reduces the amount of Foxp2-expressing cells in the developing cortex (Hawes et al. 2015). Interestingly too, core candidates for the evolution of our ability to learn and use languages (aka, our *language-readiness)* are related to vitamin D homeostasis and function (see Benítez-Burraco and Boeckx 2015 for details). At the same time, Western European Neanderthals have been claimed to suffer from vitamin D deficit, which might be related to some of their distinctive cognitive features (see Greenfied 2015 for details). Also, in present-day human populations, low vitamin D levels seem to correlate with the severity of the symptoms of cognitive diseases entailing language deficits, like autism-spectrum disorder (Bakare et al. 2011; Jia et al. 2015) and schizophrenia (Amato et al. 2010; Yüksel et al. 2014). Intriguingly, the expression of *SPAG5,* highlighted above, is regulated by VDR, the receptor of vitamin D.

We expect that this analysis in silico helps achieve a better understanding of the evolutionary changes in the regulatory landscape of *FOXP2* that seemingly contributed to refine the biological machinery supporting human speech and language. A second step will be to provide with a functional validation of our results, by mimicking the Neanderthal/Denisovan change in the binding site for SMARCC1 in the same human neuroblastoma cell line that we used for testing the functionality of the two human enhancers.

## Methods

The identification, characterization, and functional validation of the two *FOXP2* enhancers were performed as described in Torres-Ruiz et al. (2016). For the alignment of the vertebrate sequences, we used the UCSC/Penn State Bioinformatics comparative genomics alignment pipeline. The high-coverage Neanderthal and Denisovan sequences of the two enhancers were retrieved from The Neanderthal Genome Project (http://cdna.eva.mpg.de/neandertal/altai/AltaiNeandertal/bam/) and The Denisova Genome Consortium (http://cdna.eva.mpg.de/denisova/), respectively. The sequences were aligned with Clustal Omega to find the nucleotide changes. For determining the regulatory properties of the enhancers, we used the Encyclopedia of DNA Elements (ENCODE, https://genome.ucsc.edu/ENCODE/), searching for distinctive DNA hallmarks (DNase I hypersensitive sites, presence of histones with specific post-translational modifications, in particular histone H3, lysine 4 monomethylation (H3K4Me1) and H3 lysine 27 acetylation (H3K27Ac)), and particularly, for target sequences for potential co-activators and co-repressors, as revealed by chromatin immunoprecipitation followed by deep sequencing (ChIP-seq).

## References

Amato R, Pinelli M, Monticelli A, Miele G, Cocozza S (2010) Schizophrenia and vitamin D related genes could have been subject to latitude-driven adaptation. BMC Evol Biol 10: 351.

Asami M, Pilz GA, Ninkovic J, Godinho L, Schroeder T, Huttner WB, Götz M (2011) The role of Pax6 in regulating the orientation and mode of cell division of progenitors in the mouse cerebral cortex. Development 138: 5067–78.

Bakare MO, Munir KM, Kinney DK (2011) Association of hypomelanotic skin disorders with autism: links to possible etiologic role of vitamin-D levels in autism? Hypothesis 9:e2.

Benítez-Burraco A, Boeckx C (2015) Possible functional links among brain‐ and skull-related genes selected in modern humans. Front. Psychol. 6: 794.

Benítez-Burraco A, Longa VM, Lorenzo G, Uriagereka J (2008) Also Sprach Neanderthalis or did She? Biolinguistics 2.2–3: 225–232.

Coutinho P, Pavlou S, Bhatia S, Chalmers KJ, Kleinjan DA, Heyningen V (2011) Discovery and assessment of conserved Pax6 target genes and enhancers. Genome Res 21: 1349–59.

Graham SA, Fisher SE (2013) Decoding the genetics of speech and language. Curr Opin Neurobiol 23: 43–51

Green RE, Krause J, Briggs AW, Maricic T, Stenzel U, Kircher M, Patterson N, Li H, Zhai W, Fritz MH, Hansen NF, Durand EY, Malaspinas AS, Jensen JD, Marques-Bonet T, Alkan C, Prüfer K, Meyer M, Burbano HA, Good JM, Schultz R, Aximu-Petri A, Butthof A, Höber B, Höffner B, Siegemund M, Weihmann A, Nusbaum C, Lander ES, Russ C, Novod N, Affourtit J, Egholm M, Verna C, Rudan P, Brajkovic D, Kucan Z, Gusic I, Doronichev VB, Golovanova LV, Lalueza-Fox C, de la Rasilla M, Fortea J, Rosas A, Schmitz RW, Johnson PL, Eichler EE, Falush D, Birney E, Mullikin JC, Slatkin M, Nielsen R, Kelso J, Lachmann M, Reich D, Pääbo S (2010) A draft sequence of the Neandertal genome. Science 328: 710–22.

Greenfield LO (2015) Vitamin D Deficiency In Modern Humans and Neanderthals. Outskirts Press

Hawes JE, Tesic D, Whitehouse AJ, Zosky GR, Smith JT, Wyrwoll CS (2015) Maternal vitamin D deficiency alters fetal brain development in the BALB/c mouse. Behav Brain Res 286: 192–200.

Jia F, Wang B, Shan L, Xu Z, Staal WG, Du L (2015) Core symptoms of autism improved after vitamin D supplementation. Pediatrics 135: e196–8.

Johansson S (2015) Language abilities in Neanderthals. Ann Rev Ling 1: 311–332

Maricic T, Günther V, Georgiev O, Gehre S, Curlin M, Schreiweis C, Naumann R, Burbano HA, Meyer M, Lalueza-Fox C, de la Rasilla M, Rosas A, Gajovic S, Kelso J, Enard W, Schaffner W, Pääbo S (2013) A recent evolutionary change affects a regulatory element in the human FOXP2 gene. Mol Biol Evol 30: 844–852

Meyer M, Kircher M, Gansauge MT, Li H, Racimo F, Mallick S, Schraiber JG, Jay F, Prüfer K, de Filippo C, Sudmant PH, Alkan C, Fu Q, Do R, Rohland N, Tandon A, Siebauer M, Green RE, Bryc K, Briggs AW, Stenzel U, Dabney J, Shendure J, Kitzman J, Hammer MF, Shunkov MV, Derevianko AP, Patterson N, Andrés AM, Eichler EE, Slatkin M, Reich D, Kelso J, Pääbo S (2012) A high-coverage genome sequence from an archaic Denisovan individual. Science 338: 222–6.

Narayanan R, Pirouz M, Kerimoglu C, Pham L, Wagener RJ, Kiszka KA, Rosenbusch J, Seong RH, Kessel M, Fischer A, Stoykova A, Staiger JF, Tuoc T (2015) Loss of BAF (mSWI/SNF) complexes causes global transcriptional and chromatin state changes in forebrain development. Cell Rep 13:1842–54.

Ninkovic J, Steiner-Mezzadri A, Jawerka M, Akinci U, Masserdotti G, Petricca S, Fischer J, von Holst A, Beckers J, Lie CD, Petrik D, Miller E, Tang J, Wu J, Lefebvre V, Demmers J, Eisch A, Metzger D, Crabtree G, Irmler M, Poot R, Götz M (2013) The BAF complex interacts with Pax6 in adult neural progenitors to establish a neurogenic cross-regulatory transcriptional network. Cell Stem Cell 13: 403–18.

Prüfer K, Racimo F, Patterson N, Jay F, Sankararaman S, Sawyer S, Heinze A, Renaud G, Sudmant PH, de Filippo C, Li H, Mallick S, Dannemann M, Fu Q, Kircher M, Kuhlwilm M, Lachmann M, Meyer M, Ongyerth M, Siebauer M, Theunert C, Tandon A, Moorjani P, Pickrell J, Mullikin JC, Vohr SH, Green RE, Hellmann I, Johnson PL, Blanche H, Cann H, Kitzman JO, Shendure J, Eichler EE, Lein ES, Bakken TE, Golovanova LV, Doronichev VB, Shunkov MV, Derevianko AP, Viola B, Slatkin M, Reich D, Kelso J, Pääbo S (2014) The complete genome sequence of a Neanderthal from the Altai Mountains. Nature 505: 43–9.

Reuter MS, Riess A, Moog U, Briggs TA, Chandler KE, Rauch A, Stampfer M, Steindl K, Gläser D, Joset P, DDD Study, Krumbiegel M, Rabe H, Schulte-Mattler U, Bauer P, Beck-Wödl S, Kohlhase J, Reis A, Zweier C (2017) FOXP2 variants in 14 individuals with developmental speech and language disorders broaden the mutational and clinical spectrum. J Med Genet 54:64–72

Scharff C, Petri J (2011) Evo-devo, deep homology and FoxP2: implications for the evolution of speech and language. Philos Trans R Soc Lond B Biol Sci 366: 2124–40

Shriberg LD, Ballard KJ, Tomblin JB, Duffy JR, Odell KH, Williams CA (2006) Speech, prosody, and voice characteristics of a mother and daughter with a 7;13 translocation affecting FOXP2. J Speech Lang Hear Res 49: 500–525

Torres-Ruiz R, Benítez-Burraco A, Martínez-Lage M, Rodríguez-Perales S, García-Bellido P (2016) Deletion by CRISPR-Cas9 of two enhancers rearranged in a child with speech and language impairment downregulates FOXP2 in a neuronal cell line. BioRxiv http://biorxiv.org/content/early/2016/08/02/064196

Vargha-Khadem F, Gadian DG, Copp A, Mishkin M (2005) FOXP2 and the neuroanatomy of speech and language. Nat Rev Neurosci 6: 131–138

Watkins KE, Dronkers NF, Vargha-Khadem F (2002) Behavioural analysis of an inherited speech and language disorder: comparison with acquired aphasia. Brain 125: 452–464

Yüksel RN, Altunsoy N, Tikir B, Cingi Külük M, Unal K, Goka S, Aydemir C, Goka E (2014) Correlation between total vitamin D levels and psychotic psychopathology in patients with schizophrenia: therapeutic implications for add-on vitamin D augmentation. Ther Adv Psychopharmacol 4: 268–75.

